# Genome-wide changes in genetic diversity in a population of *Myotis lucifugus* affected by white-nose syndrome

**DOI:** 10.1101/764647

**Authors:** Thomas M. Lilley, Ian W. Wilson, Kenneth A. Field, DeeAnn M. Reeder, Megan E. Vodzak, Gregory G. Turner, Allen Kurta, Anna S. Blomberg, Samantha Hoff, Carl J. Herzog, Brent J. Sewall, Steve Paterson

## Abstract

Novel pathogens can cause massive declines in populations, and even extirpation of hosts. But disease can also act as a selective pressure on survivors, driving the evolution of resistance or tolerance. Bat white-nose syndrome (WNS) is a rapidly spreading wildlife disease in North America. The fungus causing the disease invades skin tissues of hibernating bats, resulting in disruption of hibernation behavior, premature energy depletion, and subsequent death. We used whole-genome sequencing to investigate changes in allele frequencies within a population of *Myotis lucifugus* in eastern North America to search for genetic resistance to WNS. Our results show low F_ST_ values within the population across time, i.e. prior to WNS (Pre-WNS) compared to the population that has survived WNS (Post-WNS). However, when dividing the population with a geographical cut-off between the states of Pennsylvania and New York, a sharp increase in values on scaffold GL429776 is evident in the Post-WNS samples. Genes present in the diverged area are associated with thermoregulation and promotion of brown fat production. Thus, although WNS may not have subjected the entire *M. lucifugus* population to selective pressure, it may have selected for specific alleles in Pennsylvania through decreased gene flow within the population. However, the persistence of remnant sub-populations in the aftermath of WNS is likely due to multiple factors in bat life history.

## INTRODUCTION

The emergence and spread of multiple infectious wildlife diseases during recent decades has had devastating consequences for biodiversity (Daszak *et al.* 2000; Smith *et al.* 2006). Unfortunately, anthropogenic threats, including human-mediated introductions and climate change, appear to be the main causes of exposure to sources of infection (Gallana *et al.* 2013; Martel *et al.* 2014; Tompkins *et al.* 2015). With shifting pathogen distributions, disease-related declines in naïve wildlife often threaten the persistence of populations. Examples include chytridiomycosis, which decimated populations of amphibians globally (Daszak *et al.* 1999; Lips *et al.* 2006), and avian malaria, which caused the sharp decline of island populations of birds (van Riper *et al.* 1986). More recently, white-nose syndrome (WNS) has been described as one of the most rapidly spreading wildlife diseases ever recorded (Blehert *et al.* 2009; Frick *et al.* 2010). Since the discovery of WNS in North America in early 2006, 13 species of bats have been diagnosed with the disease in 34 U.S. states and 7 Canadian provinces (www.whitenosesyndrome.org 2020). Genetic evidence suggests that *Pseudogymnoascus destructans*, the causative agent of WNS, was introduced to North America from Europe (Wibbelt *et al.* 2010; Ren *et al.* 2012; Lorch *et al.* 2013; Minnis and Lindner 2013; Campana *et al.* 2017), where affected species do not experience associated mortality (Puechmaille *et al.* 2011; Warnecke *et al.* 2012; Wibbelt *et al.* 2013; Zukal *et al.* 2016; Harazim *et al.* 2018).

When a disease enters a naïve host population, the initial wave of infection often causes epizootics resulting in mass mortality, which may extirpate local host populations or even cause species extinction (Daszak *et al.* 1999; De Castro and Bolker 2005; Frick *et al.* 2010). Where host extirpation does not occur, disease may instead act as a selective pressure on survivors, driving the evolution of tolerance or resistance and transforming a disease from being epizootic to being enzootic (Boots *et al.* 2009; Robinson *et al.* 2012; Karlsson *et al.* 2014). Where selective pressure is strong, this may occur through rapid changes in the distribution of genetic variants associated with disease susceptibility over short timescales (Gallana *et al.* 2013) and may be detectable for generations (Groot *et al.* 2002; Di Gaspero *et al.* 2012; Sironi *et al.* 2015; Deschamps *et al.* 2016). The detection of selective sweeps on particular genes or gene families has been proposed to confirm or exclude potential mechanisms of host susceptibility or pathogen virulence (Campbell and Tishkoff 2008). However, rapid bottlenecks (such as those caused by panzootic events) are associated with a more stochastic loss of alleles, which does not necessarily indicate selection (Luikart *et al.* 1998).

The psychrophilic (cold-growing) fungus *P. destructans* (Minnis and Lindner 2013) that causes WNS (Lorch et al. 2011) acts as an opportunistic pathogen of bats, invading the skin tissues of hibernating hosts (Cryan *et al.* 2010; Meteyer *et al.* 2012). Susceptible species, such as *Myotis lucifugus* and *M. septentrionalis*, have shown population declines greater than 90% in affected hibernacula (Frick *et al.* 2015). The infection disrupts hibernation behavior of susceptible species and leads to more frequent arousals from torpor, evaporative water loss, premature energy depletion, and death of susceptible individuals due to emaciation (Willis *et al.* 2011; Reeder *et al.* 2012; Warnecke *et al.* 2013; Verant *et al.* 2014; McGuire *et al.* 2017). Naïve infected *M. lucifugus* upregulate genes involved in immune pathways during the hibernation period (Field *et al.* 2015, 2018; Lilley *et al.* 2017). These responses are weak during torpor but are robust during the intermittent arousals (Luis and Hudson 2006; Field *et al.* 2018). Therefore, increased arousals may be attempts by the host to counter the pathogen, in addition to quenching thirst, grooming, expelling waste and possibly foraging (Willis *et al.* 2011; Brownlee-Bouboulis and Reeder 2013; Bernard and McCracken 2017), and supplementing electrolytes (Cryan *et al.* 2013).

Much of the research on disease-induced selection has focused on the major histocompatibility complex (MHC), and indeed, diseases can drive the evolution and maintenance of MHC diversity in natural host populations (Paterson *et al.* 1998; Jeffery and Bangham 2000; Teacher *et al.* 2009; Spurgin and Richardson 2010; Zeisset and Beebee 2014; Davy *et al.* 2017). Yet, factors not directly associated with interactions between host and pathogen, such as environmental conditions and competition with other species, can have a considerable influence on the manifestation of a disease (Scholthof 2007). In particular, white-nose syndrome is a prime example of a disease that is manifested when factors associated with the environment (i.e. temperature and humidity of the hibernaculum), the host (and the host’s response to infection) and the pathogen (optimum growth temperature range, suitable host) overlap, i.e. intersect within the “disease triangle” (Scholthof 2007; Turner *et al.* 2011). Therefore bat species, and populations within species, are affected differently according to hibernation behavior and prevailing micro-climate conditions in available hibernacula (Johnson *et al.* 2014; Langwig *et al.* 2015; Grieneisen *et al.* 2015). As such, in hosts challenged with an opportunistic pathogen, such as *P. destuctans*, that is capable of persistence in the environment in the absence of the host, loci not associated with MHC diversity or other immune response-associated factors (Donaldson *et al.* 2017), such as the amount of body fat prior to hibernation (Cheng *et al.* 2019), may also affect survival of species and populations. This highlights the importance of examining the genome as a whole (Sparks *et al.* 2019).

The initial impacts of WNS on six species of hibernating bats in the northeastern and midwestern USA have varied from almost complete extirpation to arrested population growth at the colony scale (Turner *et al.* 2011), leading to extensive declines at larger geographic scales (Thogmartin *et al.* 2012; Ingersoll *et al.* 2013). After the initial decline caused by WNS in *M. lucifugus*, reports have surfaced describing stabilization of colonies at smaller sizes or even increases in numbers of individuals in some areas (Langwig *et al.* 2015; Maslo *et al.* 2015; Frick *et al.* 2017; Dobony and Johnson 2018). Models parameterised with long-term data on fungal loads, infection intensity and counts of *M. lucifugus* at hibernacula have suggested development of either tolerance or resistance in these persisting populations (Frick *et al.* 2017). This is supported by reports of infected individuals not arousing from torpor as frequently as during the acute phase of the zoonosis (Lilley *et al.* 2016; Frank *et al.* 2019). Because WNS has resulted in massive population declines in some *M. lucifugus*, there is a possibility that selection could occur for alleles conferring resistance or tolerance within the standing genetic variation. Indeed, Maslo and Feffermann (2015) suggested the occurrence of evolutionary rescue, and Donaldson et al. (2017) described changes in an immunome across *M. lucifugus* populations obtained through a sequence capture method, which may be attributed to selection by WNS. Langwig et al. (2017) suggested that the initially affected *M. lucifugus* population is beginning to show signs of resistance to the pathogen. However, the selective pressures WNS has exerted on the population, which may not even be related to immune responses (e.g. Cheng et al., 2019), have only recently been described at the whole-genome level (Auteri and Knowles 2020), and high-sequencing depth studies are still lacking. Furthermore, no knowledge exists on how WNS may have affected gene flow within the previously panmictic eastern population of *M. lucifugus* (Miller-Butterworth *et al.* 2014; Wilder *et al.* 2015) and if responses differ between bats in different geographic areas.

Here, we utilize high-throughput whole-genome sequencing of the WNS-affected species *M. lucifugus*, allowing us to look at entire genomes via single nucleotide polymorphisms (SNPs). We compare genetic patterns (F_ST_ and heterozygosity) before and after the spread of WNS in the eastern North American population to gain insight into whether the disease is causing selection of major loci in surviving bats. We also examine whether the disease may have decreased gene flow within the previously panmictic population, and if bats in two geographic areas show differing signs of selection, by comparing samples collected from Pennsylvania to samples collected from New York. We hypothesize that, due to the massive population declines in *M. lucifugus* caused by WNS at a large geographic scale (Ingersoll *et al.* 2016), and the possibility that the affected population may now be beginning to stabilize or even slightly increase in size in some areas (Langwig *et al.* 2017), selection based on standing genetic variation has resulted in differentiation in one or many regions of the genome (Messer and Petrov 2013). We also hypothesize that reduced gene-flow across the once panmictic population may result in higher degrees of fixation in regions of the genomes of bats from different geographic areas.

## MATERIALS AND METHODS

### Ethics Statement

This study was carried out on bats from non-endangered species in strict accordance with recommendations in the Guide for the Care and Use of Laboratory Animals of the National Institutes of Health. All methods were approved by the Institutional Animal Care and Use Committee at Bucknell University (protocols DMR-016 and DMR-17). The bats were collected under Pennsylvania Game Commission Special Use Permit #33085, State of Michigan Scientific Collector’s Permit #1475, and New York State Department of Environmental Conservation Permit #427.

### Sample collection and DNA extraction

We conducted whole-genome sequencing on a population of *M. lucifugus* prior to, and up to 10 years after the onset of WNS using a total of 219 samples (Table 1.). For historic samples, wing tissue for sequencing was obtained from frozen specimens of known origin (Supp. Table 1.) that were collected and stored by the New York State Department of Environmental Conservation (Delmar, NY) and by D.M. Reeder at Bucknell University in Pennsylvania (PA). White-nose syndrome in bats was first observed in upstate New York (NY), at Howe’s Cave during the winter of 2005-2006 (Blehert *et al.* 2009). Our historic samples were obtained from individuals from of the eastern population of *M. lucifugus* (Miller-Butterworth *et al.* 2014; Vonhof *et al.* 2015) from New York (NY) (Miller-Butterworth *et al.* 2014; Vonhof *et al.* 2015), including individuals from the first batch of dead bats found at Hailes Cave, NY, during the winter of 2006-2007 and from animals caught in central PA in 2005 and 2006 (Fig. 1., Table 1., Supp. Table 1.). Although the bat originally described as *M. lucifugus* in North America has since been divided into five non-sister species (Morales and Carstens 2018), all of the individuals sampled in our study belong to *M. lucifugus senso stricto*, with a range extending from the east coast of North America to Alaska, and furthermore, to the same previously assigned population (Vonhof *et al.* 2015; Wilder *et al.* 2015). Previous nuclear genetic studies show that differentiation is low, and there is no evidence for any major barriers to nuclear gene flow across the eastern range of *M. lucifugus* (Vonhof *et al.* 2015; Wilder *et al.* 2015). The bats from which the historic samples in NY were collected were affected by WNS at the time of collection. However, because they were amongst the first bats to be documented with the disease in North America, we believe this set of samples is representative of the population genetic structure of *M. lucifugus* prior to WNS. Bats in PA became affected by the disease in the winter of 2008–2009 and thus had not been affected by WNS at the time of sampling in 2006-2007. Samples from PA and NY in 2006-2007 are therefore called ‘pre-WNS’ hereafter.

**Table 1.** Samples used in study.

| Sample age | Classification | Total number of<br>samples | Pennsylvania* | New York* |
| --- | --- | --- | --- | --- |
| 2006 | Pre-WNS | 25 | 25 |  |
| 2007 | Pre-WNS | 27 | 4 | 24 |
| 2015 | Post-WNS | 28 | 28 |  |
| 2016 | Post-WNS | 139 | 82 | 57 |

\* WNS first detected in New York in 2006 and PA in 2009

**Fig. 1.**
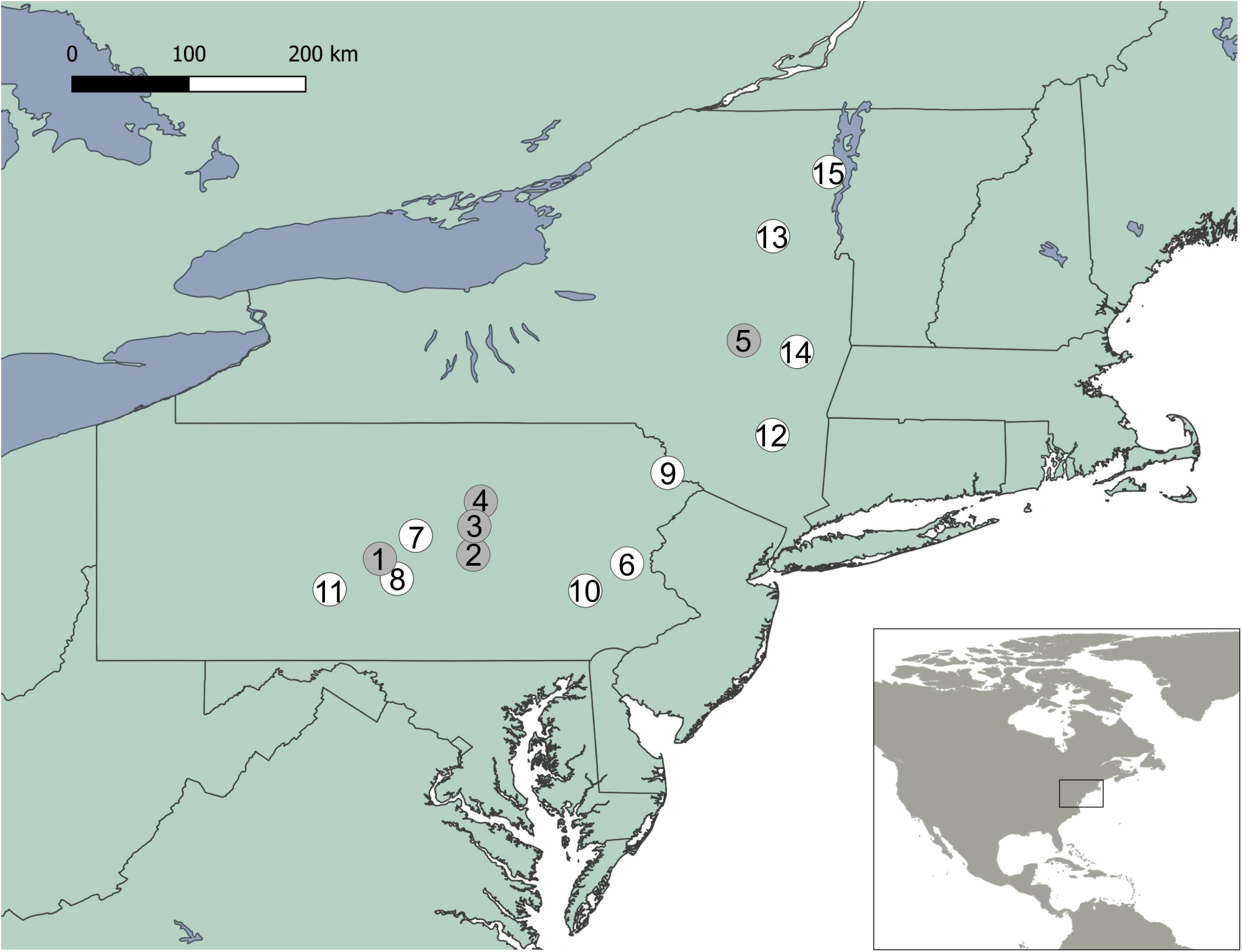
*Myotis lucifugus* sampling sites in Pennsylvania (PA) and New York (NY). Pre-WNS-sites in grey circles and Post-WNS-sites in white circles. Point 5 is Hailes Cave, near the first point of discovery of WNS. Site numbers correspond to Supplemental Table 1. Sample numbers per site 1) n=3; 2) n=19; 3) n=3; 4) n=2; 5) n=24; 6) n=32; 7) n=14; 8) n=20; 9) n=4; 10) n=12; 11) n=12; 12) n=16; 13) n=8; 14) n=36; and 15) n=21.

The remnants of the eastern North American population in NY and PA were sampled again in 2015-2016, and genetic diversity was compared before and after WNS (Fig.1., Table 1., Supp. Table 1.). Post-WNS bats were captured using mist nets and harp traps during 2015 - 2016 from a number of maternal colonies and hibernacula. In addition to samples from PA and NY, we collected gDNA from two individual bats from Upper-Peninsula Michigan (UPMI) in 2014 for use in polymorphism detection. We did this to take advantage of the greater number of polymorphic sites one would expect to arise from a more diverse collection of individuals (see sequencing methods, below). Tissue samples were collected using either 3.0 mm or 5.0 mm biopsy punches (Integra Milltex, Plainsboro, NJ) and stored in 95% ethanol until extraction. We extracted DNA using QIAmp DNA Mini Kits (Qiagen, Hilden Germany) and stored DNA at −80°C until sequencing.

### Sequencing methods

Sequencing libraries were made for two separate sets of data: **(1)** Six individuals from PA (N = 2), NY (N = 2) and UPMI (N= 2) were sequenced separately in order to initially identify a set of polymorphic sites within the *M. lucifugus* genome within the focal population (See Supp. Table 1. for information on individuals). Individual sequencing increases the confidence of SNP discovery, relative to pooled sequencing, since individuals exhibit discrete variation in the number of alleles they carry at each potential SNP site (Schlötterer *et al.* 2014); **(2)** Equimolar pools of DNA from multiple individuals from within our north-eastern US population (NY and PA) at each of the two timepoints (pre- and post-WNS) were combined to give four pooled sequencing libraries (See Table 1. and Supp. Table 1. for information on individuals). Without *a priori* knowledge of population structuring within our sampling area (although see Miller-Butterworth et al., 2014 for PA), we divided our samples in two according to geographic distance. This produced a division between the political boundaries of PA and NY. TruSeq Nano libraries with a 350 bp insert size were prepared from all samples and run on a HiSeq4000 platform to generate 2x 150 bp reads providing ∼15 Gbp of sequence data (∼7.5x coverage) for each of the six individuals **(1)** and ∼90 Gbp of sequence data (∼45x coverage) for each of the four pools **(2)**. Adapter sequences were removed with Cutadapt v1.2.1 (Martin 2011) and trimmed with Sickle 1.200 (Joshi and Fass 2011) with a minimum quality score of 20.

### Bioinformatics methods

First, reads from the six individually sequenced individuals were mapped onto the *M. lucifugus* reference genome (https://www.broadinstitute.org/brown-bat/little-brown-bat-genome-project) using Bowtie2 v2.2.5 (Langmead and Salzberg 2012), with the ‘—sensitive-local’ preset. Reads were sorted using SAMtools v1.1 (Li *et al.* 2009) and combined by read group (with duplicates removed using a ‘lenient’ validation stringency setting) using Picard-tools v2.5.0 (http://broadinstitute.github.io/picard). SNPs were called across all six individuals with the Genome Analysis Toolkit (GATK) v3.4 HaplotypeCaller (McKenna *et al.* 2010) using default parameters. Within this SNP set, high-quality bivalent SNPs were chosen that met selection criteria across the six individuals (Quality Score > 100; AF<1; 30>=DP>=100) and were input to the SAMtools mpileup function as a list of sites at which pileup was to be generated. After SNP calling, these six individuals were discarded from further analyses.

Reads from the four pooled samples were mapped to the *M. lucifugus* reference genome as above and SAMtools mpileup was used to count alleles at high-quality SNP sites identified above from the six sequenced individuals (i.e. reads matching either the reference or alternate allele for bivalent loci). Sites with coverage in the bottom 10% or top 10% quantiles summed across the pooled samples were excluded (because these could reflect inaccuracies in the reference genome), leaving approximately 13.5 million sites.

Average heterozygosity within the pooled samples was calculated using npstats (Ferretti *et al.* 2013), which controls for number of individuals within each sequencing pool and sequencing depth, thus allowing robust comparisons between pooled samples. One million SNPs were randomly picked and permuted, with 1000 sets of 1000 SNPs used to calculate the observed heterozygosity within each pooled set of individuals and, from these, we produced the median and 95% confidence intervals on heterozygosity.

To detect regions of the genome that may be under selection, we plotted F_ST_ across the *M. lucifugus* genome. Due to our sampling and sequencing regime, we were also able to separately look at individuals of the same overall population from two geographical locations within their range. In our case, an arbitrary divide was made between bats sampled in NY and PA (Figure 1), dividing our sampling population roughly at the midline. Although population genetic analysis based on a considerably lower number of genetic markers assigns *M. lucifugus* on the eastern coast of North America to the same population (Vonhof *et al.* 2015), the species shows high degrees of philopatry (Davis and Hitchcock 1965; Norquay *et al.* 2013), warranting the examination of these arbitrarily assigned pools of individuals outside the mean dispersal range of the species using more accurate, whole-genome sequencing based population genetic analyses. Using allele counts at each locus to estimate allele frequencies, F_ST_ was calculated without weighting for sampling effort 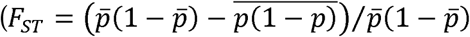, where *p* is the allele frequency and either averaged between populations 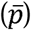 or the value *p*(1 − *p*) averaged across populations). F_ST_ was separately calculated between the Pre-WNS and Post-WNS samples and between subsections thereof, specifically PA pre- vs post-WNS, NY pre- vs post-WNS, PA pre- vs NY pre-WNS, PA post- vs NY post-WNS, and all PA vs all NY). Selection is expected to elevate F_ST_ at specific genomic regions relative to background levels due to sampling noise or genetic drift, which are expected to act equally across the genome (Holsinger and Weir 2009). To identify such regions across the genome, a moving window was applied to calculate median pairwise F_ST_ values for groups of 500 consecutive SNPs at a time to even out sampling effects. Moving windows shifted 100 SNPs at a time, thus providing the best balance between fine detail and practicality. Allele frequency estimates were weighted by the number of individuals in each pool. This was done using a custom R script (see Data availability section), run using R v3.4.1 (R Development Core Team 2011). Results were confirmed using Popoolation2 with a 10 kb sliding window and with defined pool sizes. This gave the same qualitative results, but higher levels of background noise (Kofler *et al.* 2011). Additionally, Popoolation2 was used to perform Fishers Exact Tests on individual SNPs in order to test whether any very highly significant SNPs were missed by the moving window approach.

From the comparison of the NY and PA pooled samples (i.e. the comparison that produced the highest F_ST_ values), we visually identified, across the whole genome, windows with median F_ST_ values greater than 5 standard deviations from the mean, as well as those with F_ST_ values greater than 0.05, indicating at least ‘moderate’ genetic differentiation (Wright, 1978). The numbers of such complete windows within gene models were counted for each gene to identify genes potentially under selective pressure.

## Data availability

Raw data are available at ENA accession ERP120680, and bioinformatics code at https://github.com/scottishwormboy/myoluc.

## RESULTS

### Pre-WNS vs Post-WNS comparisons across all sampled bats

Comparing F_ST_ between Pre- and Post-WNS samples revealed an overall low level of population differentiation due to genetic structure (F_ST_ median of 0.0059; Table 2). While a small number of windows have F_ST_ values slightly beyond our arbitrary cut-off of 5 standard deviations above the mean, none have F_ST_ values exceeding 0.05, suggesting that there is not even moderate selection in any region of the genome in Post-WNS samples compared to Pre-WNS samples (Fig. 2, Table 2)(Wright 1978). Similarly, using Fisher exact tests from Popoolation2 at a threshold of p < 10^−6^, no individual SNPs were found to be consistently associated with differences Pre-versus Post-WNS among the four sets of samples.

**Table 2.** Summary of individual SNP F_ST_ values across the *M. lucifugus* genome when comparing Pre-WNS with Post-WNS; and samples from PA and NY (both Pre- and Post-WNS).

|  | **Pre- vs post-WNS** | **Between site (combining pre- and post-WNS)** |
| --- | --- | --- |
| **Median (Interquartile range)** | 0.0059 (0.0013 – 0.0171) | 0.0057 (0.0013 – 0.0165) |
| **Mean (s.d.)** | 0.0127 (0.0173) | 0.0123 (0.0170) |

**Fig. 2.**
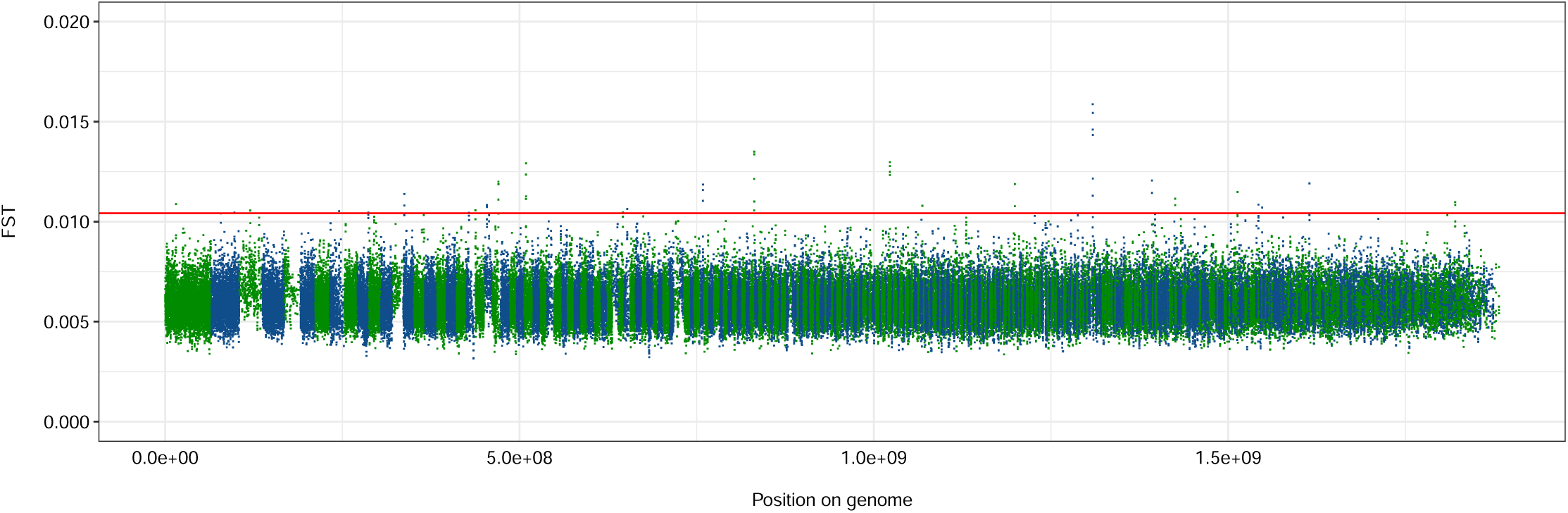
Fixation indices (F_ST_) across the *M. lucifugus* genome, comparing the population before and after the arrival of white-nose syndrome. Scaffold lengths presented in green and blue. Solid red line indicates cut-off for F_ST_ values of 5 standard deviations from the mean across the whole genome.

### Comparisons between bat populations in different geographic locations

Data split by geographic location (PA versus NY) revealed a number of minor differences (F_ST_ > 5 stdevs above mean, but F_ST_ < 0.05) by geographic location prior to arrival of WNS, as well as a peak in scaffold GL429835 with an F_ST_ value exceeding 0.05 (Fig. 3A). The high peak consists of 4 windows, the midpoints of which span an 81,887 bp region within a coding sequence consisting of 47 exons, which encodes the NIPBL cohesion loading factor (NCBI Gene ID: 102430106). The highest peak in a comparison of PA populations Pre- and Post-WNS (Fig. 3B) lies in a similar position within this coding region, as does a peak in a similar comparison of NY samples (Fig. 3C), though neither exceed the F_ST_ value cut-off of 0.05.

**Fig 3.**
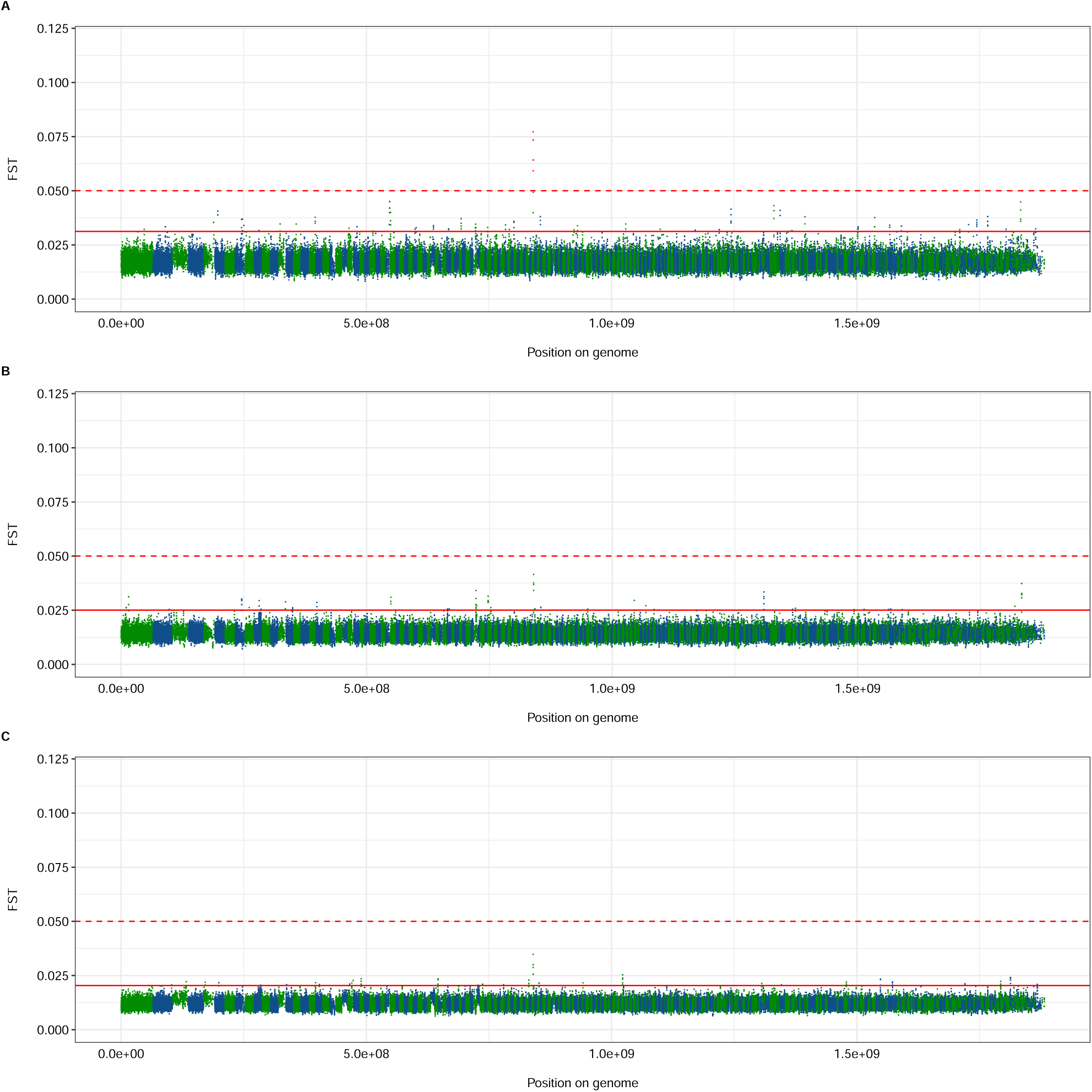
Fixation indices (F_ST_) across the *M. lucifugus* genome between geographically separated individuals of the study population: A) comparing individuals from PA to individuals from NY before the arrival of WNS; B) comparing individuals from PA before and after the arrival of WNS; and C) comparing individuals from NY before and after the arrival of WNS. Scaffold lengths presented in green and blue. Solid red line indicates cut-off for F_ST_ values of 5 standard deviations from the mean across the whole genome; dotted red line indicates cut-off of F_ST_=0.05. Windows exceeding this cutoff of F_ST_=0.05 are colored red.

When comparing the Post-WNS individuals sampled from PA with the Post-WNS individuals from NY, the peak on scaffold GL429835 is no longer visible, but a far more pronounced spike in F_ST_ values can be clearly seen in scaffold GL429776 (Fig. 4A), indicating relatively large allele frequency differences between the sample sets in this region of the genome. This is also seen, to a lesser extent, when comparing all samples from NY with all samples from PA (Fig. 4B). However, this significant peak in this comparison of all samples collected does not affect average fixation indices across the genome relative to the other comparison of all samples, Pre-WNS vs Post-WNS (Table 2). This peak could be attributed to mapping error, but our coverage depths show an average depth of 66.33x across the whole genome, whereas the focal region was sequenced at an average depth of 66.49x. This means that there has been no collapsing of duplicated regions, nor poor coverage of the region. Also, viewing the data on the NCBI genome viewer tool, with RepeatMasker histogram enabled, suggested no immediately obvious increase in repetitive sequence in the region of interest.

**Fig 4.**
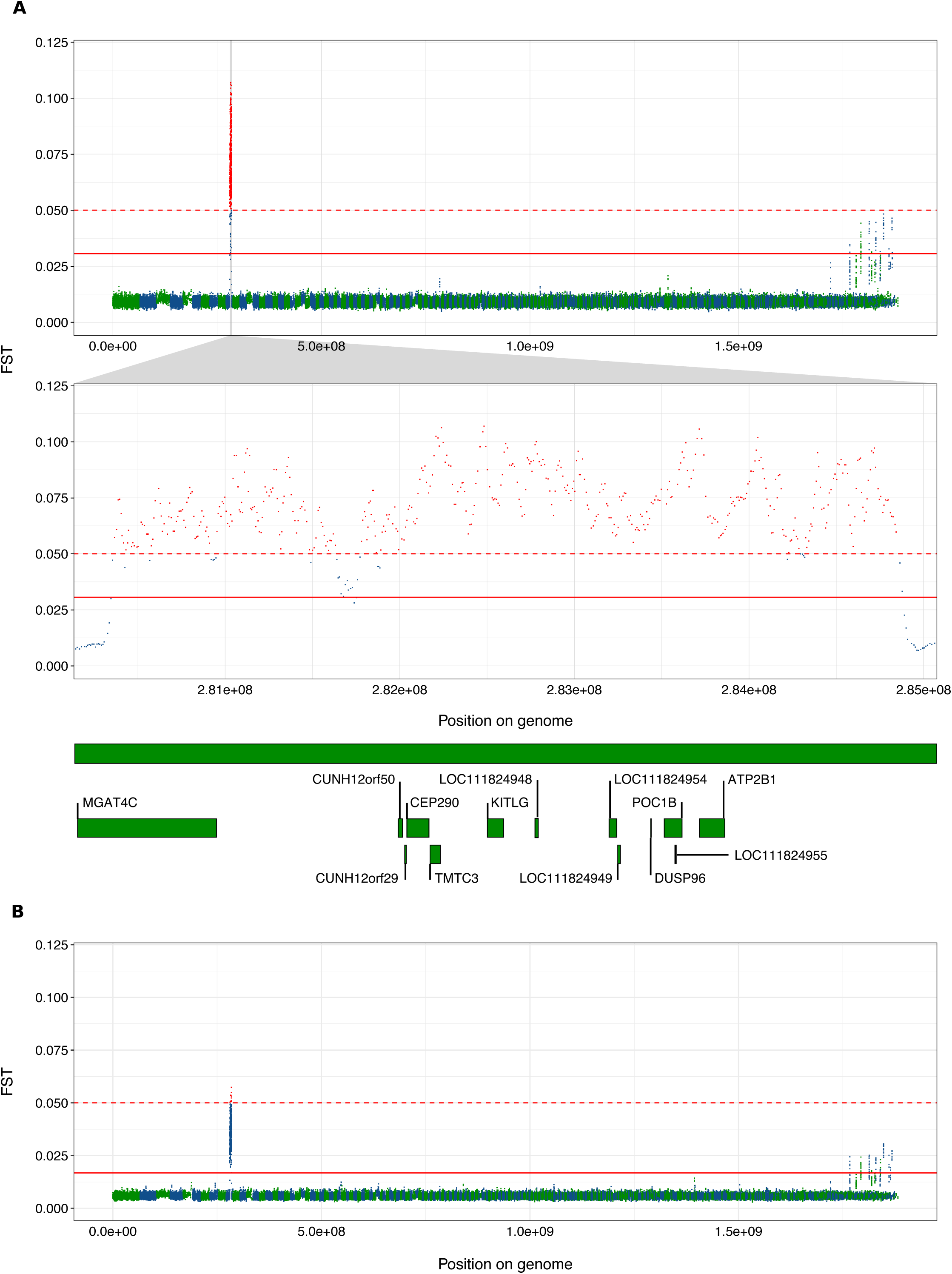
Fixation indices (F_ST_) across the *M. lucifugus* genome between geographically separated individuals of the study population: A) comparing individuals from PA to individuals from NY after the arrival of WNS, with a zoomed plot of scaffold GL429776 and gene models therein containing windows demonstrating F_ST_ values > 0.05; and B) comparing individuals from PA to individuals from NY with Pre- and Post-WNS data combined. Scaffold lengths presented in green and blue. Solid red line indicates cut-off for F_ST_ values of 5 standard deviations from the mean across the whole genome; dotted red line indicates cut-off of F_ST_=0.05. Windows exceeding this cutoff of F_ST_=0.05 are colored red.

We examined scaffold GL429776 more closely to compare the F_ST_ values between sample sets from NY Post-WNS and PA Post-WNS (Fig. 4A). Genes within scaffold GL429776 containing windows with fixation indices > 0.05 between the NY Post-WNS and PA Post-WNS samples, are presented in Table 4 and Fig. 4A. To clarify why we saw no fixation at this site when comparing Pre-WNS samples from both geographic locations (Fig. 3A), nor when we separately compared Pre-WNS and Post-WNS populations from each geographic location (Figs. 3B and 3C), we calculated the proportion of reference alleles called at each site for bats from each geographic area included in the F_ST_ window analyses (Fig. 5). This revealed a small decrease over time in the proportion of the reference alleles seen at this locus in the PA population (Fig. 5A), commensurate with an increase in proportion of reference alleles in the NY sample set (Fig. 5B). As such, existing differences between the NY and PA sample sets were insufficient to cause a relative rise in fixation indices prior to WNS spreading through the population (Fig 5C). Conversely, Post-WNS, allele proportions at this site in both sample sets had not changed sufficiently to suggest selection *within* the sample set but demonstrated clear differences *between* the NY and PA sample sets (Fig. 5D).

**Table 3.**
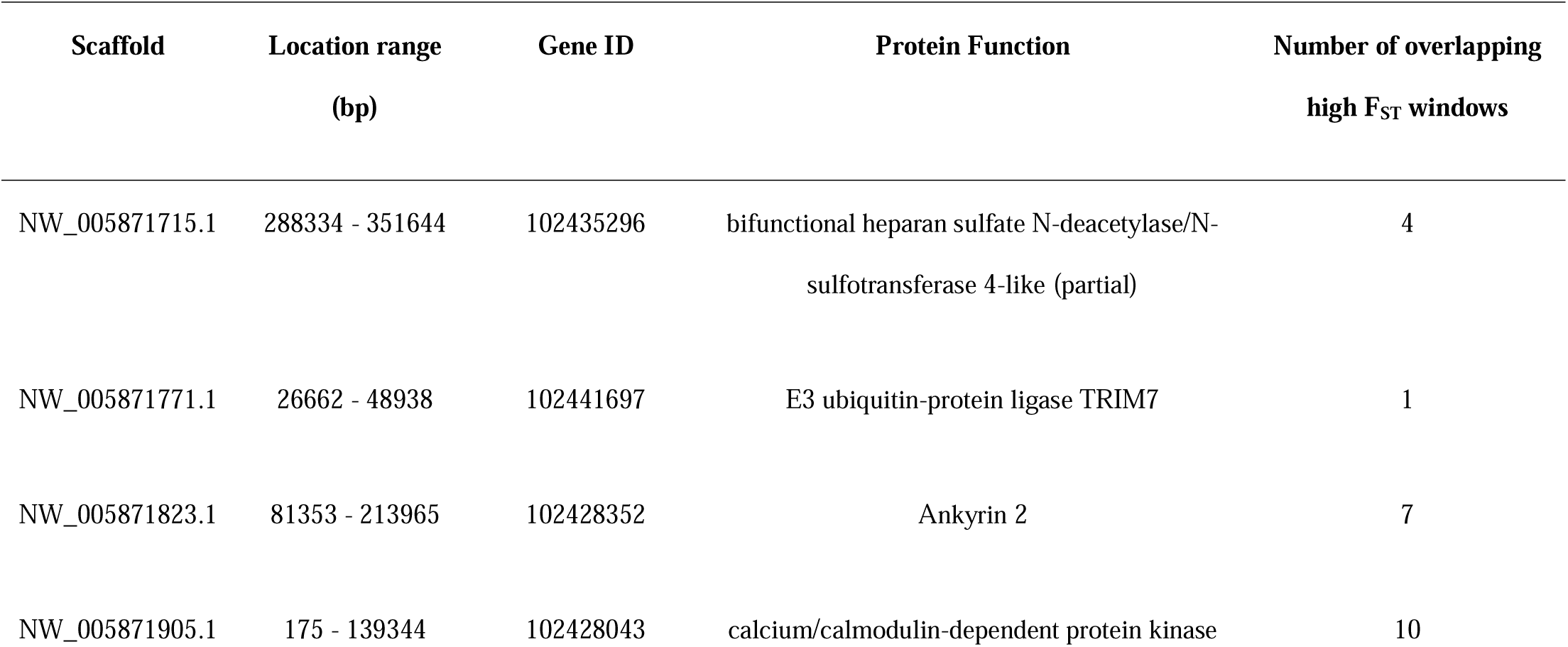
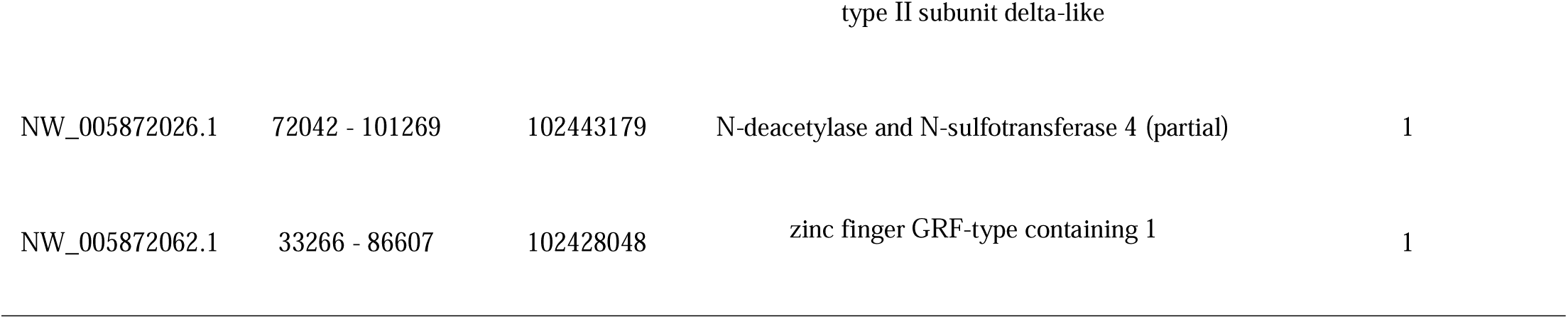
Genes within the *M. lucifugus* genome – excluding scaffold GL429776 – with windows possessing F_ST_ values >= 0.05 when comparing PA and NY samples Post-WNS.

**Table 4.**
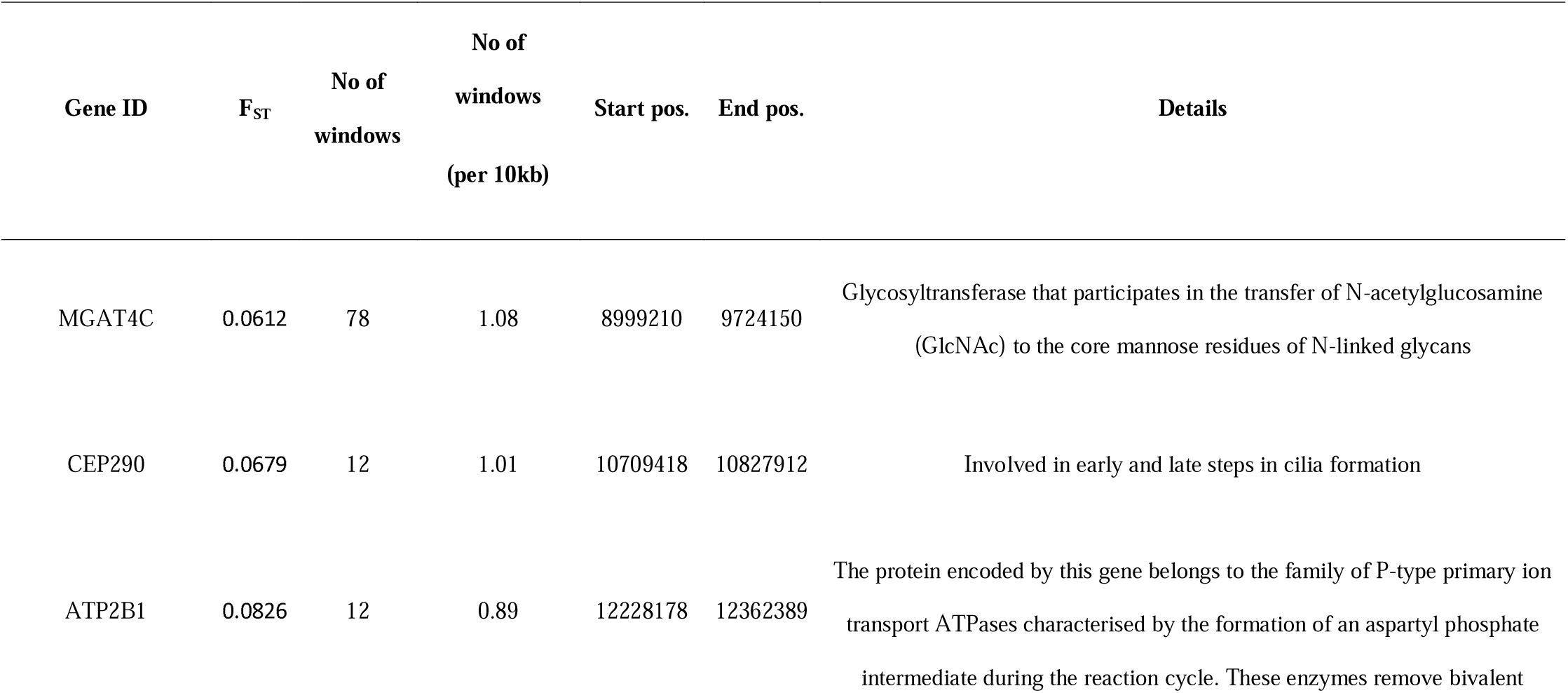
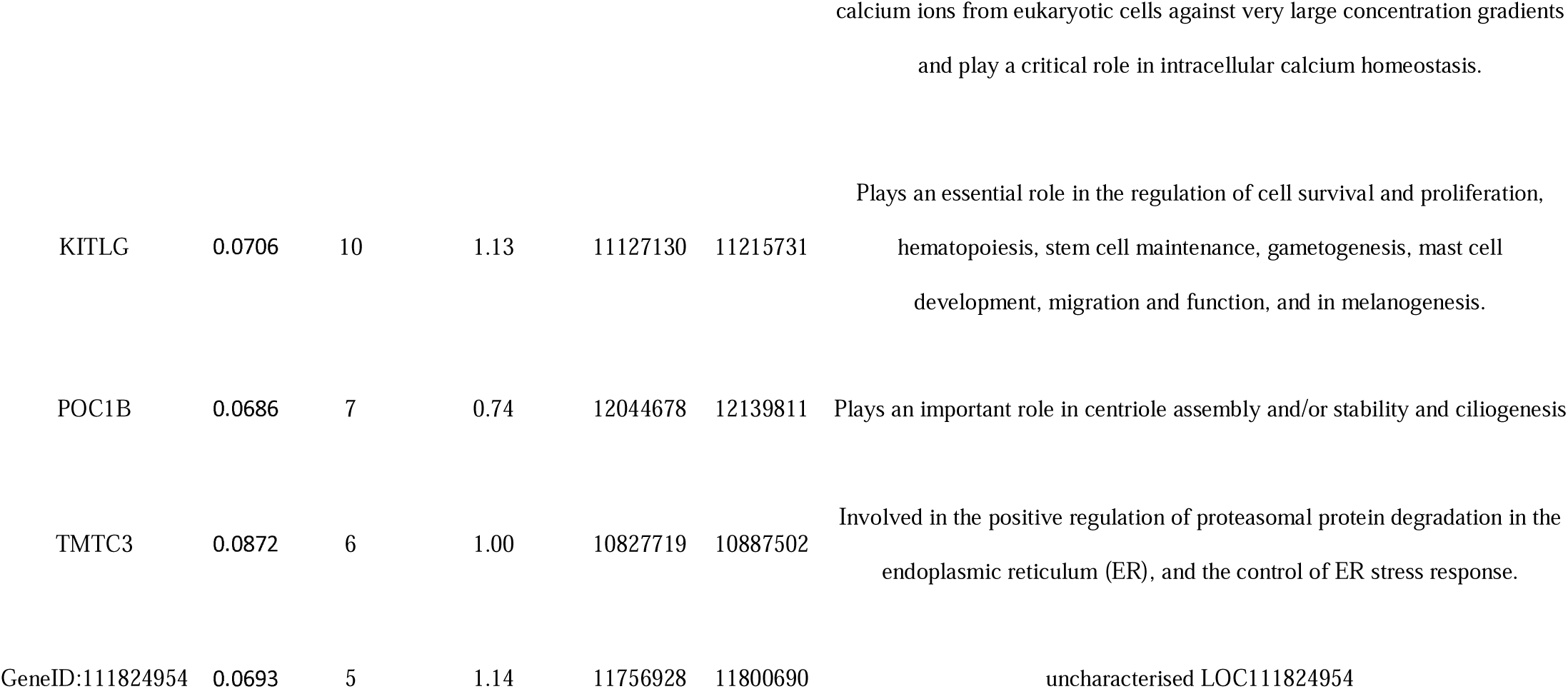
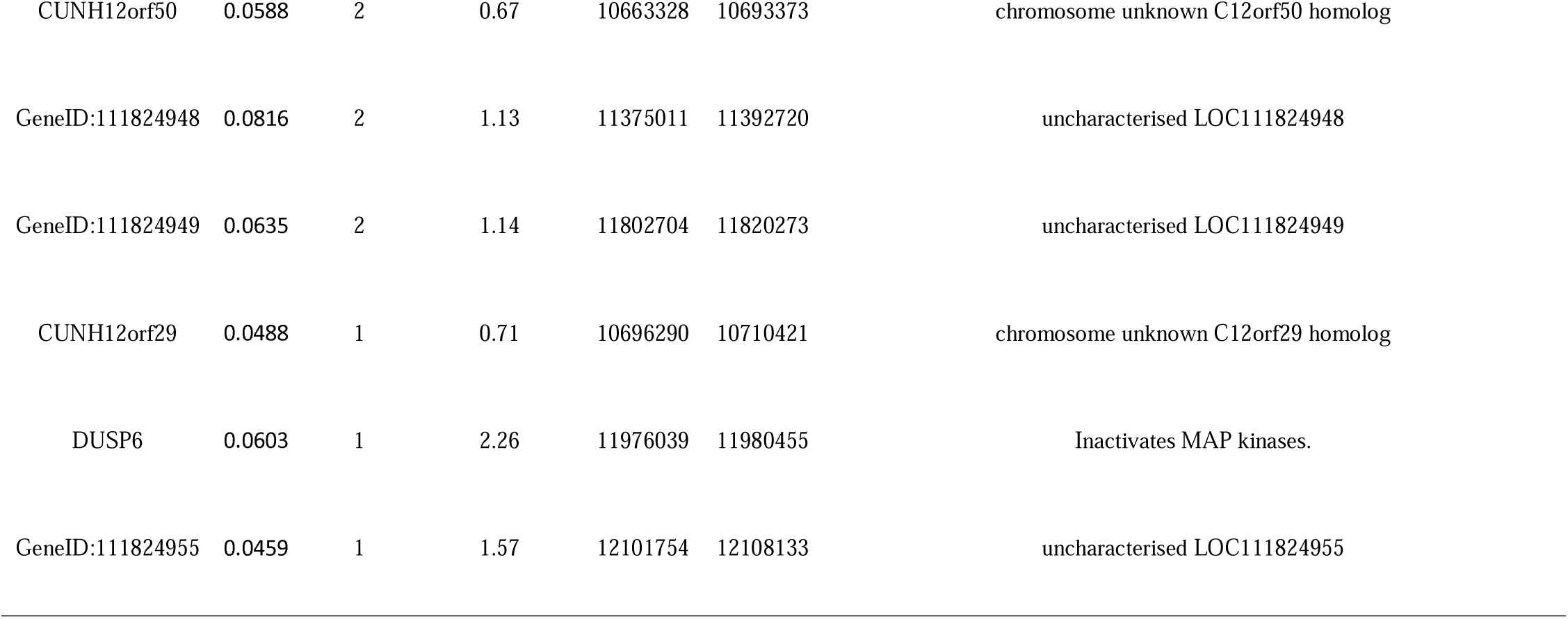
Genes within scaffold GL429776 with windows possessing F_ST_ values >= 0.05 when comparing PA and NY samples Post-WNS.

**Fig 5.**
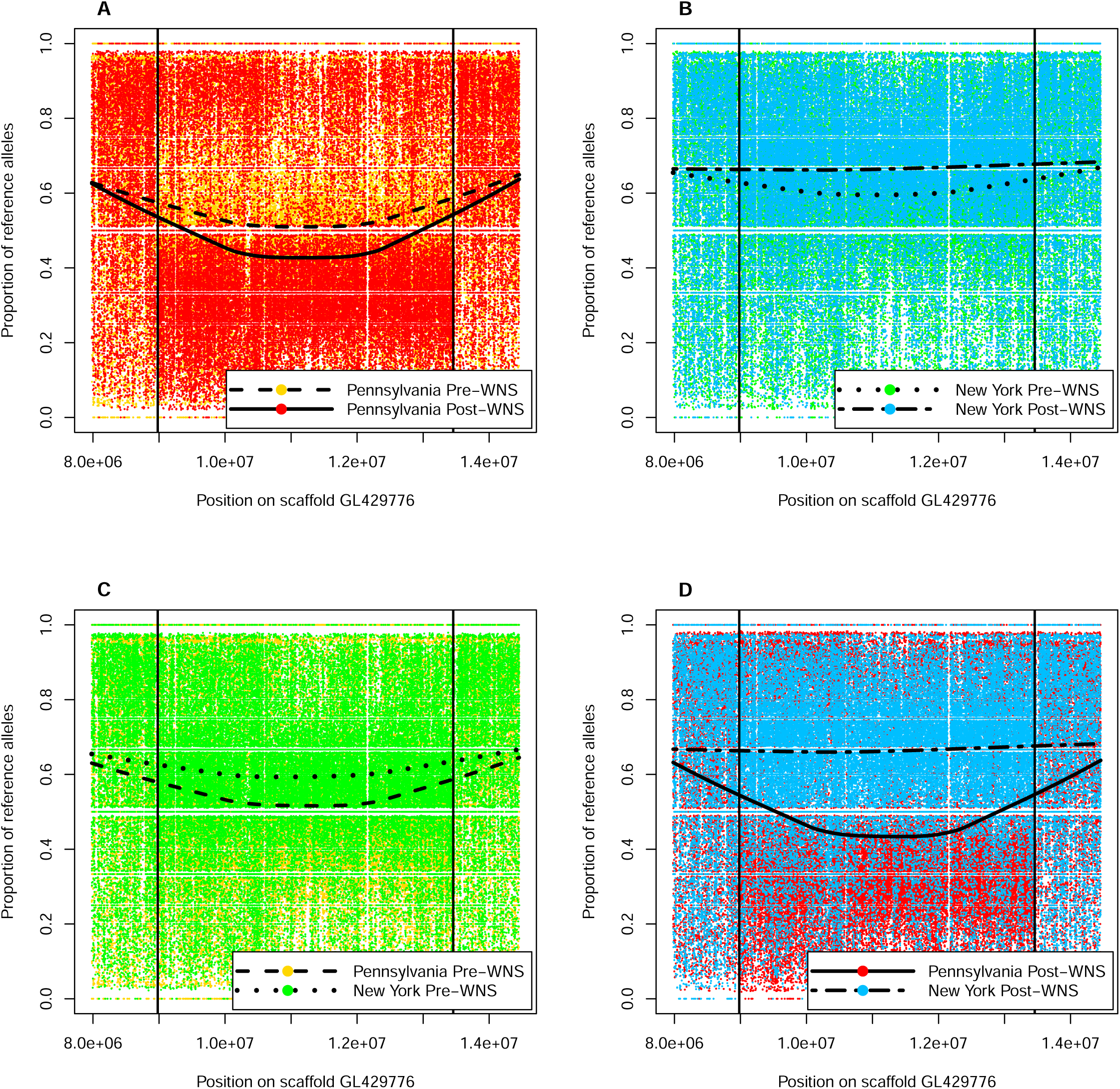
Proportions of reference alleles called at individual SNP sites for the region of scaffold GL429776 demonstrating high F_ST_ values when comparing: A) individuals from Pennsylvania before and after the arrival of WNS; B) individuals from NY before and after the arrival of WNS; C) individuals from PA to individuals from NY before the arrival of WNS; and D) individuals from PA to individuals from NY after the arrival of WNS.

In the Post-WNS comparison, peaks that were above our arbitrary threshold of 5 standard deviations above the mean across the whole genome, but below F_ST_ = 0.05, can also be observed in small scaffolds on the right-hand side of the F_ST_ plots (Fig. 4A). The 63 windows comprising these peaks lie in 47 scaffolds. Of these, high F_ST_ windows lie in only 6 protein-coding regions across 6 scaffolds (Table 3). Two of these encode partial gene models. Of the 4 remaining models containing high F_ST_ regions, all but 1 is described on NCBI as a “Low Quality Protein”. The exception is a gene encoding the ankyrin 2 protein, within which 7 high F_ST_ windows lie.

### Measures of heterozygosity within geographically and temporally-distinct sample sets

We examined heterozygosity across the genome for each geographic location at each timepoint to measure whether the epizootic reduced genetic diversity. No change in genetic diversity was detected in either geographic location: PA pre-WNS: Median: 0.263 (0.252 – 0.274 95% CI); PA post-WNS: 0.247 (0.239 – 0.255); NY pre-WNS: 0.255 (0.247 – 0.263); and NY post-WNS: 0.256 (0.247 −0.265).

## DISCUSSION

Examining our data as a whole, we found low F_ST_ values between the Pre- and Post-WNS individuals of our eastern North American *M. lucifugus* population. Also, we found no significant changes in within-population heterozygosity between Pre- and Post-WNS samples in either PA or NY, despite the massive population declines caused by WNS over the past decade, and found overall high heterozygosity across the genome. Together, these data suggest that changes in population size due to WNS have not been sufficient to affect genetic structure or diversity between time points.

Geographic separation of our bats down the middle of our total sampling region allowed us to identify a region of the genome that showed differentiation between Post-WNS PA and Post-WNS NY samples. A closer examination revealed that differentiation at this region was present already prior to WNS, but fixation in the PA sample set had increased between our two temporal sampling points. Here, we see low F_ST_ values between the two groups of bats prior to the onset of WNS, indicating an absence of selection across the genome, with the exception of a small peak in scaffold GL429835 (Fig. 3A), which lies within a gene model encoding the NIPBL cohesion loading factor. Somewhat elevated F_ST_ values are also visible within this region when comparing the PA population before and after the spread of WNS, as well as the NY population pre- and post-WNS (Figs. 3B and 3C). However, this peak is not evident when comparing PA and NY post-WNS (Fig. 4a). Any role of spatio- or temporal selection on this region is therefore weak and unlikely to be related to WNS. The region encodes the protein NIPBL, which plays an important role in the function of the cohesin complex, which is vital for chromosome segregation, DNA damage repair, and gene expression regulation (Peters *et al.* 2008). What role variation in this region could play in natural populations is unknown.

Our overall results confirm that the pre-WNS bats from NY were genetically similar to pre-WNS bats from PA. We should note, however, that it is possible that our NY samples collected from dead bats at Hailes Cave in February 2007 (Site 5, Fig. 1, Supp. Table. 1), a year after WNS was first detected at a single site in NY (Blehert et al., 2009), may not fully represent the genetic composition of the Pre-WNS NY bat population and that our analysis could have benefitted from the sequencing of samples from a variety of sites. Regarding Post-WNS sampling; because individual maternal colonies and hibernacula may show a higher degree of relatedness between individuals (Kurta and Murray 2002; Burns *et al.* 2014; Olivera-Hyde *et al.* 2019), we included bats from a number of these sites in the study. However, summer colonies and overwintering sites may be separated by considerable distances and therefore there will always be some degree of gene flow at the geographical scales considered, i.e., among sampling sites within a state (Norquay *et al.* 2013).

The F_ST_ comparison of Post-WNS individuals sampled from PA and NY depicts considerable genetic differentiation within scaffold GL429776 (Fig. 4A); differentiation which did not exist in the Pre-WNS comparison of individuals sampled from these geographical areas. This is a difference that would have gone unnoticed had we not made the comparison between bats sampled in NY and PA, or used fewer genetic markers instead of a whole-genome approach (Miller-Butterworth *et al.* 2014; Vonhof *et al.* 2015). The appearance post-WNS of this region of greater genetic differentiation within scaffold GL429776 when considering all geographical locations together is not, however, reflected when examining temporal changes in either geographic location on their own (Fig. 3B and Fig. 3C). A comparison of allele frequencies of individual SNPs within this region shows that individuals sampled from PA showed an increase in deviation from the reference genome over time, while a greater proportion of reference alleles were seen in NY samples Post-WNS compared with Pre-WNS. The differences in proportions, growing as they were over time, were insufficient to exhibit a high fixation index Pre-WNS, but were notably different after the disease had spread through the population. One could argue that this is due to sampling at maternal colonies and swarming sites, which has known effects on allele frequency differences (Johnson *et al.* 2015). However, we sampled at a number of maternal colonies, which would dilute this effect, and furthermore, recent studies suggest philopatry within maternal colonies may not be as high as previously expected (Olivera-Hyde et al., 2019, although on *M. septentrionalis*). Therefore, we assume our sampling of multiple maternal colonies and hibernation sites has removed any effects of relatedness within our sample sets.

The region within the scaffold GL429776 containing the peak has six annotated genes with very different functions (The UniProt Consortium 2019). Two, POC1B and CEP290, are involved in cilia development and maintenance. The ATP2B1 gene encodes a protein which belongs to the family of P-type primary ion transport ATPases, whereas TMTC3 is involved in the control of endoplasmic reticulum stress response. MGAT4C is a glycosyltransferase that participates in the transfer of N-acetylglucosamine (GlcNAc) to the core mannose residues of N-linked glycans. Finally, KITLG produces a protein involved in mast cell development, migration and function, and melanogenesis. KITLG has been documented as a target of evolution in recent studies and in humans it has been linked with thermoregulation (Yang *et al.* 2018), which is a critical component of bat life history (Studier and O’Farrell 1972). In fact, upregulation of KITLG at low temperature helps promote the production of brown fat for heat generation (Huang *et al.* 2014). Bats utilise brown adipose tissue in arousals to normothermia from different degrees of torpor (Thomas *et al.* 1990). There is also evidence of parallel evolution of pigmentation in sticklebacks and humans linked to KITLG (Miller *et al.* 2007), as well as its role in determining resistance/susceptibility to swine respiratory diseases in Erhulian pigs (Huang *et al.* 2017). KITLG, by influencing mast cell development, could also play a role in either protective or pathological immune responses to *P. destructans*, which may involve IgE-mediated recognition of secreted fungal proteins (Reeder *et al.* 2017; Field *et al.* 2018). As such, temporal changes we found in allele frequencies at this locus could, for instance, be linked to differences in hibernation site temperatures, and/or thermoregulatory needs in bats sampled at different locations (Thomas *et al.* 1990; Humphries *et al.* 2002). Spatial patterns, as seen here between the arbitrarily pooled samples from PA and NY, could be due to local adaptation, in which strong selective sweeps may be linked to gene variants favored in local interactions (Hansen *et al.* 2012; Schoville *et al.* 2012; Kyle *et al.* 2014; Rico *et al.* 2015). We lack data concerning migration patterns of the bats studied here, as well as the environmental conditions within their microfugia. Future work would benefit from these data in a bid to understand whether bats repeatedly frequent habitats with similar conditions and whether there is a significant difference in conditions across these eastern states that explains the rapid rise in allelic differentiation, and thus F_ST_ values, at this locus.

In contrast to our study, in which we did not detect evidence for an evolutionary rescue effect at a large geographic scale, Auteri & Knowles (2020)and Donaldson et al. (2017) found putative selectively driven genetic changes in local populations of *M. lucifucus*, which could have the potential to result in an evolutionary rescue from WNS. Differences in results may have arisen among the studies because of the more limited sampling or lack of repeated sampling before and after WNS in the other two studies, or because of the different geographic areas sampled. Further, the Donaldson et al. (2017) study presented subtle immunogenetic variation across a wide geographic area, with the post-WNS population sampled in the first or second winter of WNS exposure. Thus, differences among populations in that study could represent local adaptation, independent of the effects of WNS. An alternate possibility that could explain the findings of all three studies is that gene flow in the formerly panmictic eastern population of *M. lucifugus* could have been reduced following massive population declines, and adaptive responses to WNS could be beginning to emerge independently in localized areas, even as broad-scale adaptation is not evident across the range of the species. Further clarification of the role of the genes identified in each of the three studies could be revealed by studying differences in transcriptomes, infection, and survival in response to WNS during infection trials in hibernation experiments (Field *et al.* 2018; Lilley *et al.* 2019).

Despite not seeing an overarching genetic signal associated with resistance in *M. lucifugus*, many remnant populations appear to be surviving year after year and in some cases even increasing in size (Langwig et al., 2017; Lilley et al., 2016). The pathogen loads appears to be significantly lower for remnant populations compared to that of bats during the epidemic phase, during which massive declines were, which would suggest the development of resistance in these populations (Bernard et al., 2017; Langwig et al., 2017; but see Lilley et al., 2019). However, improved responses to WNS may occur through other means than the evolution of resistance. For instance, during the hibernation period, when the bat hosts are most vulnerable to the pathogen, roost microclimate variables, including humidity and temperature, affect the growth of the fungal pathogen and the survival of the host (Verant *et al.* 2012; Johnson *et al.* 2014; Grieneisen *et al.* 2015; Marroquin *et al.* 2017), and behavioral shifts in roost site selection by bat hosts since the onset of WNS (Johnson *et al.* 2016) could favor energy conservation while reducing fungal growth and infection. Modelling also suggests that environmental factors, including the cave microbiome, have an impact on the proliferation and infectivity of *P. destructans* (Hayman, Pulliam, Marshall, Cryan, & Webb, 2016; Lilley et al., 2018). This is supported by the discovery of microbes in hibernacula environments and on bats that are able to retard the growth of the fungus (Zhang *et al.* 2014, 2015; Micalizzi *et al.* 2017). In light of our results and considering bat life history as a whole, adaptation and evolutionary rescue may not be a fast track for recovery of bat populations affected by WNS (Maslo and Fefferman 2015), although ultimately these will contribute to long term survival of populations (Lilley *et al.* 2019). At the present, therefore, it is more likely that behavioral shifts in selection of hibernation sites that vary in environmental conditions and strong selection for microbial taxa that inhibit *P. destructans* could explain why some colonies have persisted or may have even begun to recover from the zoonosis (Cheng *et al.* 2016; Johnson *et al.* 2016; Lemieux-Labonté *et al.* 2017; Lilley *et al.* 2018).

More broadly, it is still unclear how frequently genetic adaptation occurs in natural populations and under what circumstances it is promoted (Schoville *et al.* 2012). Many studies have recorded phenotypic changes in natural populations and attributed them to host-pathogen interactions (Kilpatrick 2006; Råberg *et al.* 2009; Frank *et al.* 2014; Langwig *et al.* 2017). However, it is difficult to determine conclusively whether changes in phenotype are the result of selection in the genome or a result of phenotypic plasticity (Paterson *et al.* 2010; Routtu and Ebert 2015). Also, the assumed benefit of change (i.e. adaptation) is often not tested experimentally, which would allow for inference of causality with the pathogen through exclusion of other potential drivers.

To conclude, our results suggest WNS has not yet decidedly subjected populations of *M. lucifugus* to selective pressures leading to genetic adaptation in remnant populations. No population -wide signs of selection in comparisons of genomes of Pre-WNS and ten years Post-WNS populations were observed in this study (although see Donaldson *et al.* 2017; Auteri and Knowles 2020). This is something that is not unexpected with a species in which the effective population size most likely numbered in tens of millions of individuals prior to the onset of WNS. The disease has not caused a true population bottleneck, as exemplified by our measures of genetic diversity not varying between pre- and post-WNS samples. Our results indicate that the persistence and recent growth of some remnant populations of *M. lucifugus* are more likely attributable to other factors, such as microbiome adaptation and hibernation site selection among others, rather than genetic adaption. However, we found increased variability in a specific area of the genome in a comparison of Post-WNS samples from our two different geographic locations, relative to Pre-WNS samples. We suggest this is due to weakened connectivity between bats at different locations allowing for local adaptations to appear in the absence of gene flow. Although genes in the high F_ST_ region of the genome were identified, and genes such as KITLG provide interesting avenues of research, even with regards to WNS, further investigation into the processes in bat life history to which these genes are related are required to expound upon the existing gene annotations.

## ACKNOWLEDGEMENTS

We thank Svenska Kulturfonden, H2020 Marie Skłodowska□Curie Actions (TML, 706196) and the Natural Environment Research Council, UK for funding the study; staff at the Centre for Genomic Research, University of Liverpool, UK for DNA sequencing expertise; Jenni Prokkola and Riley F. Bernard for proofreading the manuscript; and Alice Lilley, Cali Wilson, Beth Rogers, Mike Scafini, Scott Wasilko, Cassandra Ostroski, Heather Rogers, Emily Blackman, Kristin Gaines, Rachel Mosely, Angela Remeika, Imran Ejotre, Laura Kurpiers, Joseph S. Johnson, Marianne Gagnon, Spencer Schell, Sarah Bouboulis, Karen El Chaar and Allentown Parks for assisting in sample collection.

